# Tracking the Brain’s Intrinsic Connectivity Networks in EEG

**DOI:** 10.1101/2021.06.18.449078

**Authors:** Saurabh Bhaskar Shaw, Margaret C. McKinnon, Jennifer J. Heisz, Amabilis H. Harrison, John F. Connolly, Suzanna Becker

## Abstract

Functional magnetic resonance imaging (fMRI) has identified dysfunctional network dynamics underlying a number of psychopathologies, including post-traumatic stress disorder, depression and schizophrenia. There is tremendous potential for the development of network-based clinical biomarkers to better characterize these disorders. However, to realize this potential requires the ability to track brain networks using a more affordable imaging modality, such as Electroencephalography (EEG). Here we present a novel analysis pipeline capable of tracking brain networks from EEG alone, after training on supervisory signals derived from data simultaneously recorded in EEG and fMRI, while people engaged in various cognitive tasks. EEG-based features were then used to classify three cognitively-relevant brain networks with up to 75% accuracy. These findings could lead to affordable and non-invasive methods to objectively diagnose brain disorders involving dysfunctional network dynamics, and to track and even predict treatment responses.

## 1. Introduction

A large body of neuroimaging research over the past decade indicates that the brain is organized into functional networks of interacting brain regions, called intrinsic connectivity networks (ICNs). The study of large-scale ICNs has provided considerable insight into the neural basis of human cognition and behaviour in the healthy and diseased brain [52, 23]. There is immense potential to use features of ICN dynamics as clinical biomarkers in patients with various psychopathologies [23] and even track their response to treatments. However, the predominant imaging modality used to study ICNs, functional magnetic resonance imaging (fMRI), is expensive, lacks the necessary temporal resolution, and is not readily available for performing routine neurocognitive assessments in patients with brain disorders. A major breakthrough would be to track ICNs using a non-invasive and more widely accessible modality such as electroencephalography (EEG). Here we present a significant advancement towards this goal. We apply machine learning methods to simultaneously recorded EEG and fMRI data, to derive supervisory signals for learning EEG-based ICN features, permitting highly accurate classification and tracking of ICNs from EEG alone.

Three predominant networks have been extensively studied [23] using fMRI: the central executive network (CEN) generally involved in exteroceptive processing, *i.e*. tasks involving attention to external stimuli; the default mode network (DMN) involved in interoceptive processing tasks, *e.g*. autobiographical memory retrieval, imagining the future, spatial planning and navigation, and self-reflection; and the salience network (SN), thought to modulate switching between CEN-mediated exteroceptive and DMN-mediated interoceptive cognitive processes [44]. Appropriate network switching dynamics between these three core networks is thought to be critical for healthy cognitive functioning. Disruptions in normal inter- and intra-network connectivity in these ICNs have been observed in numerous neuropsychological conditions affecting emotion and cognition. For instance, those with Major Depressive Disorder (MDD) [7] exhibit deficits in downregulating activity within the DMN in association with persistent rumination [29]. Those with Post-Traumatic Stress Disorder (PTSD) exhibit disruptions in the activation and functional connectivity of DMN, CEN and SN [2, 34, 20, 48]. For example, abnormal activation of the DMN has been observed in PTSD patients while switching to a working memory task that normally recruits the CEN [8]. Disrupted ICN dynamics are also observed in numerous other psychological disorders [23], such as bipolar disorder [56, 51], schizophrenia [57] and mild cognitive impairment (MCI) [55]. Therefore, it is extremely promising to track the network dynamics within these ICNs as clinical markers of brain disorders. The ability to monitor such network activity will be particularly useful for tracking progress in the treatment of such neuropathologies, where network dynamics is dysregulated, and may also lead to the development of novel individualized treatments such as network-based neurofeedback interventions.

Unfortunately, doing so using fMRI would be prohibitively expensive. Moreover, fMRI lacks the temporal resolution to track the temporal dynamics of the networks on a millisecond timescale. An appealing alternative is electroencephalography (EEG), a cheaper and more widely accessible imaging modality with excellent temporal resolution.

In previous work some EEG features of these brain networks have been identified. For instance, working memory load on the CEN is found to modulate frontoparietal EEG power in theta and upper alpha frequency bands [37], fronto-parietal phase-based functional connectivity graphs [10], and common spatial patterns [5]. Frontal lobe EEG activity in the theta frequency band is also found to negatively correlate with the DMN [39], and when combined with delta and alpha band powers, is capable of discriminating the DMN from the sensorimotor network [41]. Furthermore, theta-gamma coupling is a key mechanism driving hippocampal memory processes required by the DMN during autobiographical memory retrieval and is found to be dysfunctional in patients with memory impairments [28]. Hence, a combination of within-frequency and cross-frequency phase-based and amplitude-based connectivity measures could capture various component neural processes inherent to the CEN, DMN and SN. An open question is whether unique signatures of each of these networks can be identified, allowing us to track each of the three networks as distinct from the other two.

Rather than focusing on such temporal dynamics, most previous attempts to identify EEG features of ICNs have relied on spatial filtering analyses such as beamforming and blind source separation [46]. These analyses are especially susceptible to signal leakage due to volume conduction [32], and specific data acquisition parameters [21], limiting their utility as clinical tools for tracking ICN dynamics. One alternate approach is to track EEG microstates [17], which are spatial correlates of ICNs identified by spatial clustering. However, despite its increasing popularity in probing dysfunctional ICN dynamics in numerous psychological conditions [26], this analysis is riddled with flawed assumptions that lead to inaccuracies at finer temporal scales [43, 27].

To date no one has identified unique signatures that permit classification and tracking of ICNs using EEG alone. To accomplish this, we developed a machine learning model that learns EEG signatures of the ICNs using supervisory labels derived from simultaneously recorded fMRI. The simultaneous EEG-fMRI data were collected from two cohorts of participants that performed two different multi-task paradigms: a dual task-switching paradigm designed to activate the DMN, CEN and SN [44]; and a multi-task paradigm that cycles through a series of seven tasks, a subset of which rely on the three networks of interest [16]. A large battery of approximately 40M amplitude and phase-based features were computed from the EEG data collected during these tasks. Classification labels for the DMN, CEN and SN were derived from the thresholded activity of these ICNs, identified using group-wide ICA analysis of the simultaneously acquired fMRI data. These labels were used to select the optimal feature set for a multiclass SVM classifier, using a hierarchical minumum-redundancy-maximum-relevance (mRMR) [33] feature selection algorithm. Importantly, having identified these features from the simultaneous EEG-fMRI data, there is the potential to track the ICNs using EEG data alone. The ability of the EEG features alone to classify the predominantly active network was validated on testing sets from the above described datasets using 20-fold crossvalidation.

## 2. Results

### 2.1. FMRI Networks as labels

The first step in our analysis pipeline (see Figure 1C) was to identify the “ground-truth” ICN activity, *i.e*. the correct network labels, as identified using fMRI. This involved a group-wide independent component analysis (group-ICA) to discover components that overlap with the ICNs of interest.

**Figure 1:**
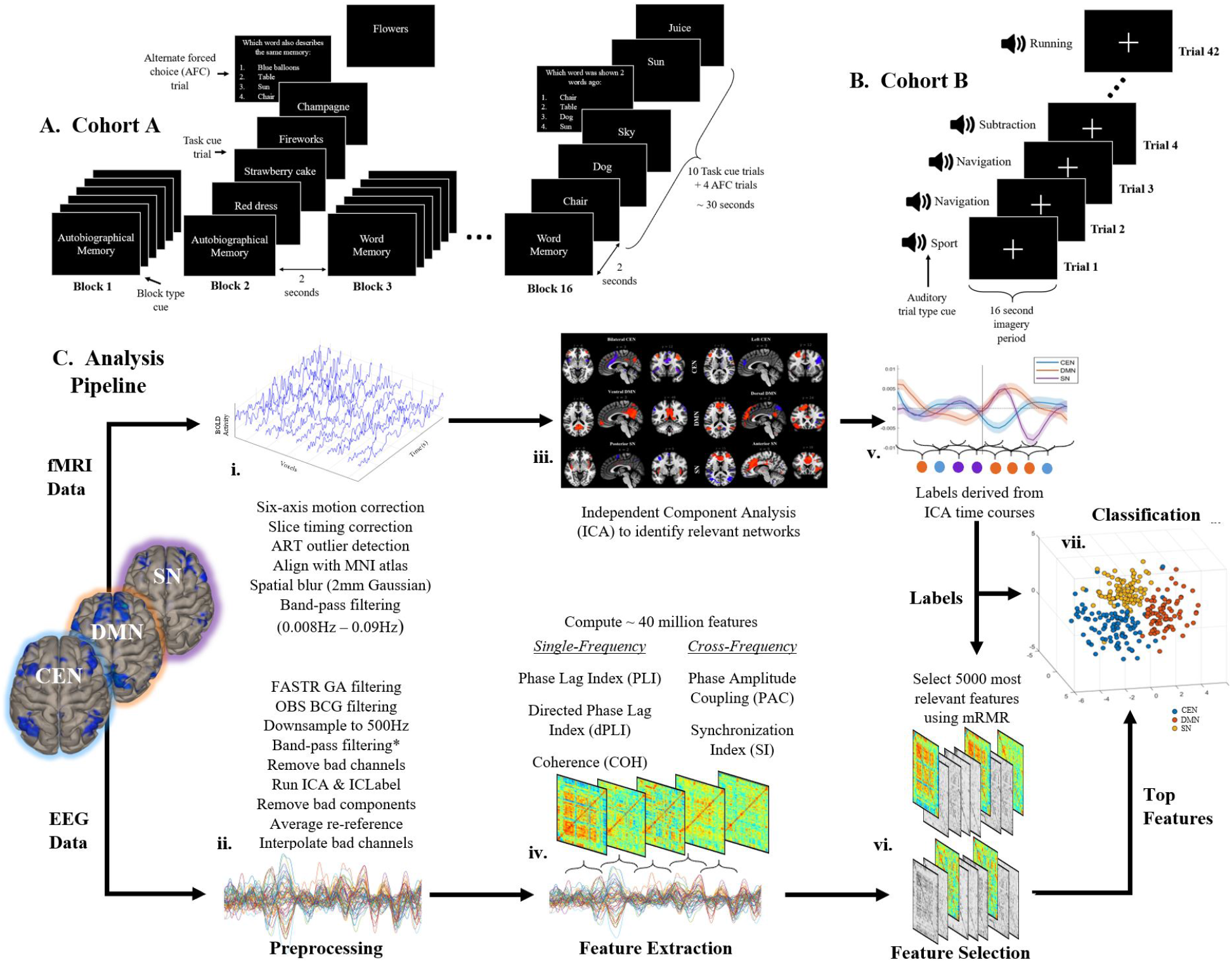
The simultaneous EEG-fMRI data used in this study were collected from two cohorts of participants that performed two different multitask paradigms, shown in panels A and B. Cohort A used a dual task-switching paradigm (shown in panel A) designed to activate the DMN, CEN and SN, identical to the paradigm used in Shaw *et al*. [44]; and Cohort B performed a multi-task paradigm (shown in panel B) that cycled through a series of seven tasks [16], a subset of which relied on the three networks of interest. Refer to the methods section for a detailed description of the tasks. Panel C details the analysis pipeline used in this study to analyze the simultaneous EEG-fMRI data collected from Cohorts A and B. The top row illustrates the analysis steps for fMRI data (sub-panels i., iii., and v.), while that for EEG data is shown in the bottom row (sub-panels ii., iv., and vi.). *Top row* (*fMRI data analysis*): i. Preprocessing - The fMRI scans were first realigned and unwarped, followed by motion correction, slice timing correction (STC), ART-based outlier identification and scrubbing, normalization to the MNI152 atlas, spatial smoothing using a 2mm Gaussian kernel, and band-pass filtering between 0.008Hz–0.09Hz.; iii. The fMRI data was then decomposed into 20 independent components using ICA, of which 6 components were found to be relevant to the DMN, CEN and SN (shown here). v. Classification labels for the DMN, CEN and SN were derived from the thresholded activity of these three ICNs. *Bottom row*(*EEG data analysis*): ii. The concurrently collected EEG data was preprocessed by first removing the gradient artefact (GA) using a parallel optimized version of the FASTR gradient artefact removal toolbox Shaw [42], followed by ballistocardiogram (BCG) filtering using optimal basis set filtering (OBS). The artefact-free EEG data were then downsampled from 5000Hz to 500Hz, followed by temporal band-pass filtering into six different frequency bands - full (1-50Hz), delta (1-4Hz), theta (4-8Hz), alpha (8-13Hz), beta (13-30Hz), and gamma (30-50Hz). Bad channels were then detected and removed based on their spectral characteristics, followed by an ICA decomposition to identify and remove artefacutal components such as ocular artefacts, eye blinks and muscle artefacts. Finally, the EEG data were referenced to average EEG channel data, after which the previously removed bad channels were interpolated using spherical interpolation. iv. A large battery of approximately 40M amplitude and phase-based features were computed from the preprocessed EEG data. vi. The previously derived classification labels were used to select the optimal feature set for a multiclass SVM classifier, using a hierarchical minumum-redundancy-maximum-relevance (mRMR) [33] feature selection algorithm. vii. These features were used to classify the three ICNs. The clear separation of the three classes shown results in high classification accuracy.

**Figure 2:**
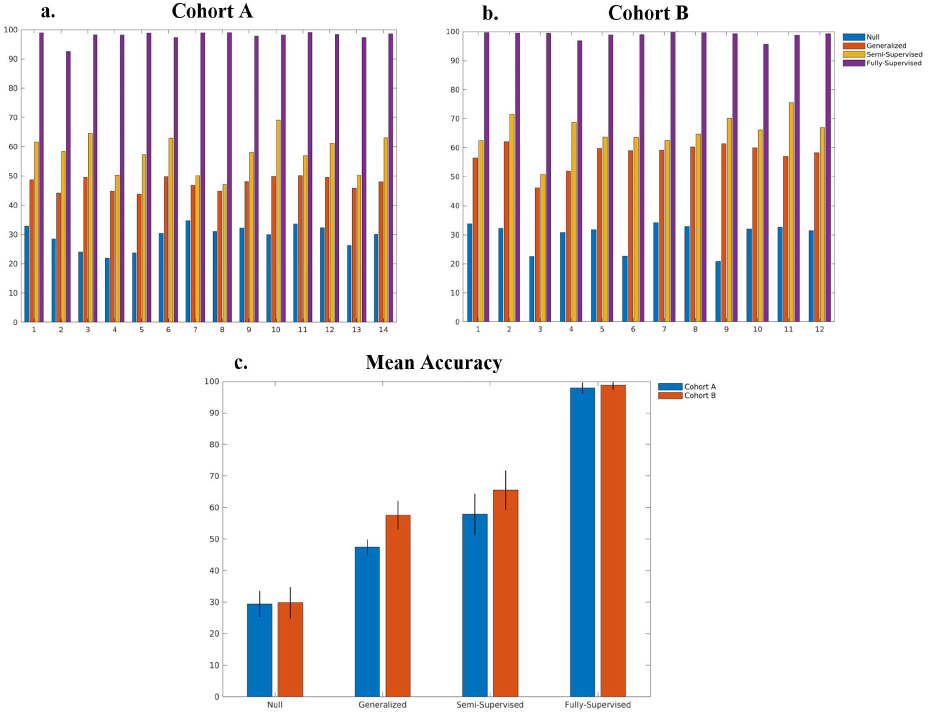
The test-set classification accuracies for predicting the activation of CEN, DMN and SN networks using EEG data alone across cohorts A and B, shown in (a.) and (b.) respectively. The accuracy of the null model is shown in blue, that of the generalized model is shown in orange, and that of the fully-supervised model is shown in yellow. while that of the surrogate null model, trained using random labels, is shown in red. The average classification accuracy for the three classifiers is shown in (c.), with cohort A shown in blue and cohort B shown in orange.

The group-ICA analysis identified a total of 20 components with distinct spatio-temporal patterns of activity. Of these, 11 components showed significant spatial overlap (high Dice coefficient) with well known ICNs [45], while the remaining 9 components represented unwanted noise and artefacts. We focused on the 6 components representing CEN, DMN and SN subnetworks (Figure 1C.iii), and extracted the time-courses of overall ICN activity by averaging the activity of each ICNs’ component sub-networks (Figure 1C.v). The temporal dynamics of these components matched those expected from the CEN, DMN and SN within the dynamic task-switching paradigm used in cohort A, with the SN causally influencing the CEN and DMN in a task-linked manner. The fMRI temporal dynamics of the ICNs lie outside the scope of this paper and are further explored in Shaw *et al*. [44].

The component time courses for the CEN, DMN and SN were partitioned into overlapping windows of 5 fMRI time points, using a sliding-window approach and advancing by 1 fMRI time point for each window. Each time window was labeled with the most active ICN during the corresponding time period, creating the class labels for feature selection and classification of the EEG data to predict the predominantly active ICN during each time window.

### 2.2. Learning EEG features from fMRI-derived labels

The next step in our processing pipeline (see figure 1C) involved using the fMRI labels identified in section 2.1 to select relevant EEG features.

#### 2.2.1. Generalized EEG features

A generalized feature set was identified using mRMR on the features extracted from the EEG data, using labels derived from the simultaneously acquired fMRI data. Here mRMR tries to find EEG features that are maximally relevant to the fMRI-derived class labels across all participants, while being minimally redundant.

#### 2.2.2. Individualized EEG features

An individualized EEG feature set was also identified using a similar procedure for each individual participant. Here mRMR tries to find separate EEG features for each participant that are maximally relevant to the fMRI-derived class labels for that participant, while being minimally redundant.

Having obtained these features, they can now be applied to EEG data from other datasets. We now investigate the ability of these features to predict ICN activity. Researchers can then utilize the trained classifiers with the corresponding EEG features dervied from their own datasets to assess ICN activity using EEG data only.

### 2.3. ICN activation can be identified using EEG features alone

Using the optimal feature sets identified in the previous section, we trained three versions of the classifier to predict CEN, DMN and SN activation, representing three scenarios of data availability while using this pipeline. The first two scenarios (generalized classifier and semi-supervised individualized classifier) represent a use case where the researcher has access to only EEG data from their participants. They can use our models trained on EEG-fMRI derived features to predict ICN activity using only EEG data from their participants. The third scenario (fully-supervised individualized classifier) represents a use case where the researcher has access to simultaneous EEG-fMRI data from their own participants. The observed classification performance for each of the three scenarios is detailed below.

#### 2.3.1. Generalized classifer

The first classifier was trained on the generalized feature set from all but one participant. This classifier was then tested on the left-out participant’s EEG data, comparing the predicted labels to the “ground-truth” labels derived from their corresponding fMRI data. These classifiers achieved an average classification accuracy of 58% ± 6% for cohort A and 61% ± 5% for cohort B, and performed significantly better than the corresponding surrogate null models (p < 0.001), trained using randomly permuted training labels.

#### 2.3.2. Semi-supervised individualized classifier

We then explored a semi-supervised approach to boost the classification performance of the generalized classifier, by training a custom classifier for each participant’s EEG data using the predicted class labels from section 2.3.1. To accomplish this, the time points with a confidently predicted label (maximum posterior probability > 75%) from the generalized classifier were picked as the labelled time points, while the rest of the time points were considered unlabelled. When comparing these EEG-only derived labels were then compared to the “ground-truth” labels from the participants’ fMRI data. the confidently labeled time points boosted the average classification accuracy to 82% ± 6%, while accounting for only 25% of the participant’s time points. Therefore, this process split each participants’ data into a small labelled dataset with “expert” labels from the generalized classifier, and a larger unlabelled dataset.

Optimal features were selected using a weighted-mRMR approach that used labels predicted by the generalized classifier, and a weighted mutual information estimate for identifying minimally redundant and maximally relevant features. The weighted mutual information estimate weighted each time point by the maximum posterior probability of the predicted label, giving more importance to the time points with more confident predictions. A semi-supervised approach was then used to predict the labels of the unlabelled data, achieving an average accuracy of 57.9% ± 6% for cohort A and 65.5% ± 6% for cohort B.

#### 2.3.3. Fully-supervised individualized classifier

Lastly, a fully-supervised individual classifier was trained on a subset of each participants’ data, using the individualized feature set identified in section 2.2.2 with their fMRI-derived labels. This classifier was tested using a held-out subset of the same participants’ data, achieving an average classification accuracy of 98% ± 3% for cohort A and 96% ± 4% for cohort B.

Interestingly, the expansion of the EEG signal from 64 channels to an extremely high dimensional feature space (40M), and its subsequent reduction to a 5000 dimensional space, made the classification task much easier by transforming the EEG feature space from an inseparable domain to a readily separable domain, shown in Figure1C.vii. This allowed a simple classifier, a multi-class support vector machine (SVM) to achieve the highest classification accuracy, substantially outperforming several much more complicated deep neural network classifiers (see supplementary figure). These deep neural network methods perform notoriously poorly in domains where the number of observations is not significantly larger than the number of features, where they are susceptible to overfitting.

### 2.4. Characterizing the features of each network

MRMR feature selection identified the top 5000 most relevant features for discriminating between the CEN, DMN and SN. To gain a more intuitive understanding of the EEG signatures uniquely representative of each ICN’s activity, a Shapley additive explanations (SHAP) analysis [22] was conducted for each feature within the identified feature space for the generalized classifier. The SHAP analysis explains the contribution of each feature to the classification, by identifying the relative change in log-odds of each ICN label, given an increase in the corresponding feature value. The SHAP values of each feature for the correct ICN label, averaged across all trials, are shown in Figure 3 for the single-frequency features and in Figure 4 for the cross-frequency features. These are further discussed in the following sections.

**Figure 3:**
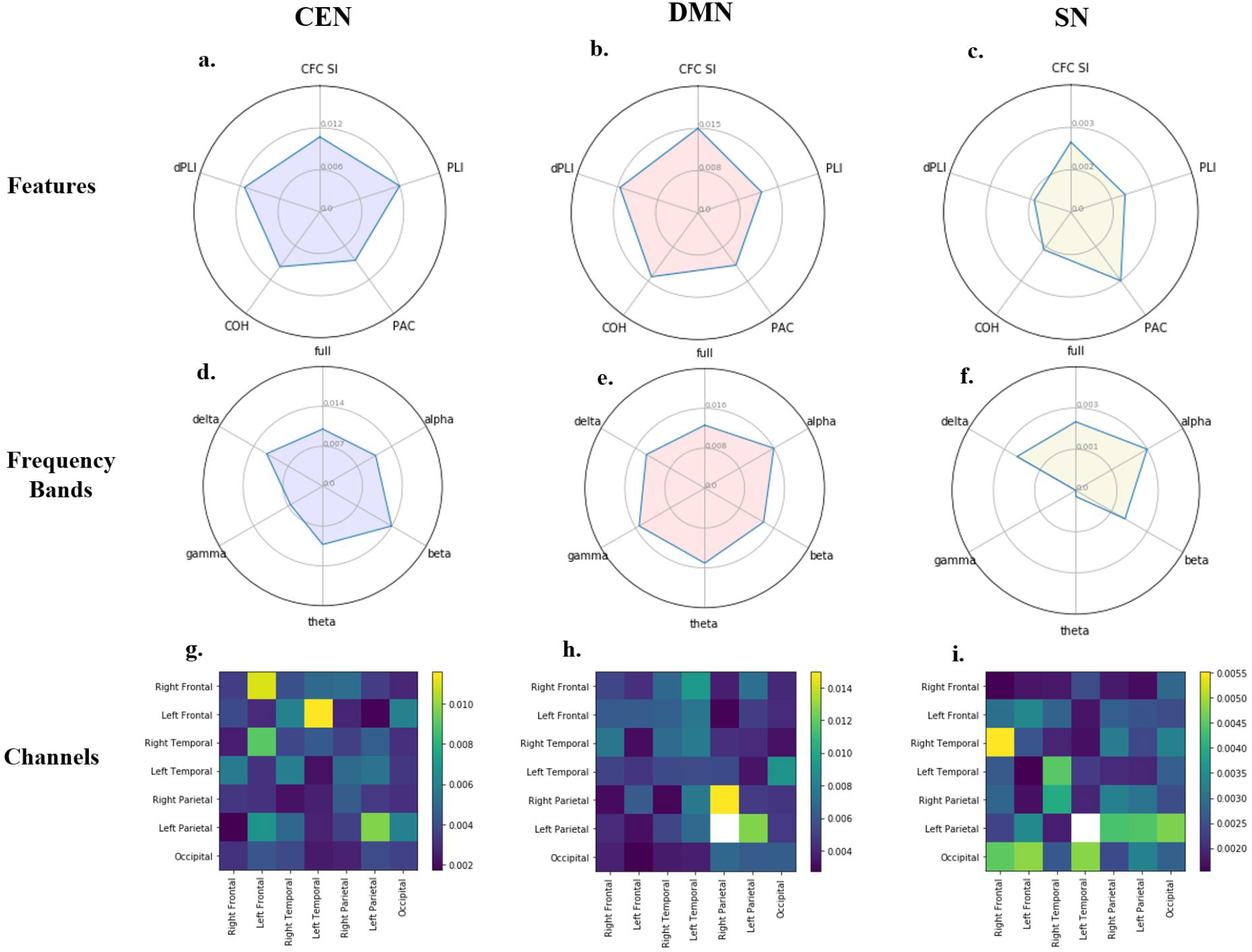
The influence of the identified feature set in increasing the log-odds of each network class (CEN, DMN and SN). The contributions of the five features across different frequency bands and channel pairs are shown in the top, middle and bottom rows respectively.

**Figure 4:**
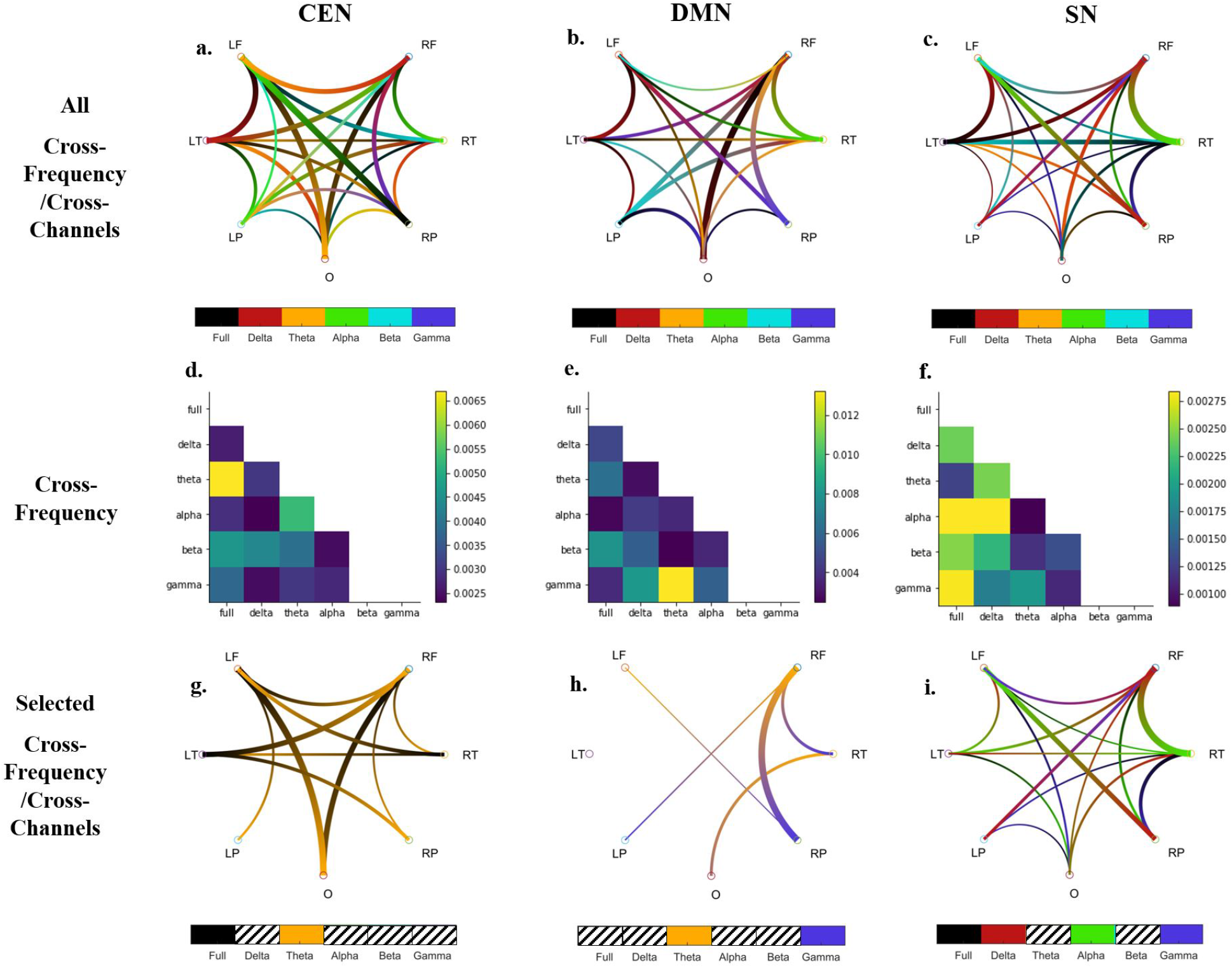
The influence of the identified feature set in increasing the log-odds of each network class (CEN, DMN and SN), shown for the crossfrequency features (SI and PAC).

Five connectivity features were included in our feature set - three single frequency features, and two cross frequency features. The single frequency features identified the connectivity between pairs of EEG channels within the same frequency band, and included phase lag index (PLI), directed phase lag index (dPLI), and coherence (COH). The cross-frequency features describe the connectivity between pairs of EEG channels across different frequency bands, and include phase amplitude coupling (PAC) and synchronization index (SI).

### 2.5. CEN Features

All five connectivity features contributed towards the classification of the CEN network, with the PLI and dPLI being the highest contributors (Figure 3a). Among the single-frequency features, high connectivity between fronto-temporal, intra-parietal and intra-frontal electrodes (Figure 3g) across theta, alpha, beta and delta bands (Figure 3d) were found to be indicative of CEN activity.

**Table 1:** Three-way classification accuracy (test set) for predicting the activation of the CEN, DMN and SN across cohort A and B. The reported values are the cross-validated (20-fold) mean accuracy of the multiclass-SVM model trained using a 75-25 train-test split of the data. All classification accuracies of the trained model are significantly higher than that of the surrogate null model, trained using random labels.

| Pt.<br>Num-<br>ber | Cohort A |  |  |  | Pt.<br>Num-<br>ber | Cohort B |  |  |  |
| --- | --- | --- | --- | --- | --- | --- | --- | --- | --- |
|  | Null<br>Class. | Gen.<br>Class. | Semi-sup.<br>Ind.<br>Class. | Fully-sup.<br>Ind.<br>Class. |  | Null<br>Class. | Gen.<br>Class. | Semi-sup.<br>Ind.<br>Class. | Fully-sup.<br>Ind.<br>Class. |
| 1 | 32.9 | 48.7 | 61.6 | 98.9 | 1 | 33.8 | 56.4 | 62.4 | 99.7 |
| 2 | 28.5 | 44.2 | 58.4 | 92.6 | 2 | 32.3 | 62.1 | 71.5 | 99.4 |
| 3 | 24.0 | 49.5 | 64.6 | 98.2 | 3 | 22.6 | 46.2 | 50.8 | 99.4 |
| 4 | 22.0 | 44.8 | 50.3 | 98.1 | 4 | 30.8 | 51.9 | 68.8 | 96.9 |
| 5 | 23.7 | 43.8 | 57.4 | 98.8 | 5 | 31.7 | 59.7 | 63.6 | 98.8 |
| 6 | 30.4 | 49.8 | 62.9 | 97.3 | 6 | 22.7 | 59.0 | 63.5 | 98.9 |
| 7 | 34.7 | 46.9 | 50.1 | 98.9 | 7 | 34.2 | 59.1 | 62.5 | 99.8 |
| 8 | 31.0 | 44.8 | 47.2 | 99.0 | 8 | 32.9 | 60.2 | 64.7 | 99.6 |
| 9 | 32.3 | 48.1 | 58.1 | 97.8 | 9 | 20.8 | 61.4 | 70.1 | 99.2 |
| 10 | 30.0 | 49.9 | 69.1 | 98.1 | 10 | 32.0 | 59.9 | 66.1 | 95.6 |
| 11 | 33.6 | 50.2 | 57.0 | 99.1 | 11 | 32.7 | 57.1 | 75.6 | 98.7 |
| 12 | 32.4 | 49.5 | 61.2 | 98.3 | 12 | 31.4 | 58.3 | 66.9 | 99.2 |
| 13 | 26.3 | 45.8 | 50.3 | 97.3 | - | - | - | - | - |
| 14 | 30.1 | 48.0 | 63.0 | 98.5 | - | - | - | - | - |

Interestingly, the features most strongly influencing the CEN classification showed two unique lateralization patterns (Figure 3g). The fronto-temporal and intra-parietal connectivity features were left-lateralized, alongside bilateral and cross-hemispheric fronto-temporal and intra-frontal connectivity features. These two lateralization patterns are consistent with the bilateral and left-lateralized sub-networks of the CEN, seen with the fMRI results in Figure 1C.iii.

A wide range of cross-frequency coupling (CFC) features were also found to influence CEN classification, as shown in Figure 4a. Among these, the connectivity features between theta band and the full frequency band were particularly predictive of CEN activity (Figure 4d). Furthermore, the channel-pairs communicating within these frequency bands included interhemispheric frontal-frontal and fronto-temporal channels pairs, along with intra-hemispheric fronto-parietal and fronto-occipital channels. The direction of the fronto-temporal connections followed a theta to full-band direction, with the phase of the theta frequency at the frontal electrodes synchronizing with the full frequency band activity at the temporal electrodes. This pattern is consistent with frontal theta-driven processes during executive tasks such as working memory [38] and mental arithmetic [36].

### 2.6. DMN Features

Similar to the pattern seen in CEN classification, all five connectivity features were informative in predicting DMN activity, with dPLI, COH and SI being more informative than PLI and PAC. However, in contrast to the pattern seen in CEN classification, the most informative single-frequency features included gamma, theta and alpha band activity within parietal-parietal channel pairs. Furthermore, the intra-parietal connectivity within the single-frequency features was restricted to ipsi-lateral channel pairs with minimal inter-hemispheric connections. Such parietal-driven gamma activity could be indicative of self-related processing occurring within key parietal DMN nodes such as the posterior cingulate cortex (PCC) and the Precuneus.

Additionally, fronto-parietal theta-gamma coupling features were found to be the most informative CFC feature, following a fronto-parietal direction, with the theta phase of frontal electrode activity synchronizing with the gamma activity of the parietal electrodes. These included bilateral cross-hemispheric fronto-parietal connections, and a right-lateralized fronto-parietal connection, with the latter being most informative of DMN activity. These data are consistent with a large body of research implicating theta-gamma coupling in the various DMN-linked memory processes [12, 35, 31].

### 2.7. SN Features

In contrast to the pattern seen for CEN and DMN classification, the cross-frequency coupling features (SI and PAC) dominated the prediction of SN activity. The single-frequency features that contributed to SN classification included a wide range of frontotemporal, temporal-temporal, occipito-frontal, parietal-parietal and parieto-occipital connections. Furthermore, these connections were within the beta, alpha, delta and full frequency band, with no single-frequency features within the gamma and theta bands.

Among the CFC features, coupling between alpha-delta, gamma-full band and alpha-full band was found to be most predictive of SN activity, across a wide combination of channel-pairs. This is consistent with the integrative role of the SN that involves communication with numerous brain regions across a wide range of frequency bands.

One notable observation is the concentration of gamma and delta activity at the right frontal electrodes, that might be linked to the activity of the right anterior insula (rAI) node of the SN. This node is particularly important for task-linked switching between the CEN and DMN, and is well connected with other frontal, parietal and temporal brain regions [47, 25, 24].

Taken together, the identified EEG feature set captures critical spatio-temporal characteristics of the CEN, DMN and SN, that is consistent with their functional roles and previously observed dynamics.

## 3. Discussion

In this study, we used cutting-edge machine learning methods to classify and track the activity of three major ICNs using EEG data, which was previously only possible using fMRI data. We focused on three core ICNs within Menon’s tri-network model [23] - the CEN, DMN and SN, given their relevance in characterizing a wide range of psychopathologies. FMRI studies indicate that these intrinsic networks are dysregulated in psychopathologies including PTSD, and can even be used to predict patients’ PTSD subtype [30]. Moreover, characterizing ICN dynamics has the potential to track response to treatments and inform individualized treatment planning decisions. However, despite its potential as a clinical assessment tool, probing ICN dynamics using fMRI is prohibitively expensive. Other barriers to routine clinicial use of fMRI in psychiatric disorders include the distress caused by the confined enclosure and loud sounds made by the scanner, both of which can be triggering for those with PTSD. EEG, by comparison, is much more affordable, relatively portable, and does not carry with it the risk of claustrophobia- and noise-related distress. Therefore, the potential to monitor brain networks using EEG, as afforded by the present study, can greatly improve the clinical accessibility of ICN-based assessment.

To accomplish this, we used fMRI-derived ICN labels to select relevant features from simultaneously-acquired EEG data and classified the active ICNs using EEG data alone. Three distinct classification scenarios were explored, providing investigators with three levels of performance, depending on the type of data available to them. The first scenario involves applying the generalized model trained on our dataset to EEG data from other participants to predict the active ICN. The second scenario extends this approach by additionally applying semi-supervised learning to train a custom classifier for each participant, improving classification accuracy by 5% to 15%. These two approaches represent the most significant contribution of this work, allowing investigators to probe ICN activity using the proposed methods with EEG data from their participants. Lastly, the third scenario demonstrates that higher classification accuracy is achievable by fully-supervised learning, if the investigator has access to simultaneously acquired EEG-fMRI data from their participants.

It is important to note that the performance of the classifiers in the three scenarios was not significantly different between the two cohorts (p=0.6), demonstrating that our methods can be used to discriminate between the activity of the CEN, DMN and SN, across a wide range of cognitive tasks.

This study also investigated the EEG features contributing to the detection of each ICN, identifying EEG signatures that uniquely characterize the CEN, DMN and SN. Interestingly, the data-driven approach used in this study identified EEG signatures that aligned with major findings in the literature.

Frontal theta band activity, found to be predictive of both CEN and DMN activation in this study (Figure 4), has been implicated in both executive processes (CEN functioning) and autobiographical memory encoding and retrieval processes (DMN functioning). In the context of executive tasks, frontal theta is thought to synchronize the pre-frontal cortex with a wide range of other brain regions [14], signalling the need for cognitive control processes during periods of high risk and/or high levels of uncertainty [4]. Doing so in a phase-linked manner, it acts as an alarm signal for instantiating cognitive control processes to better learn from the higher error rates encountered during such scenarios. Adaptations in theta band dynamics also assist with optimizing this process, as observed with the reduction in peak theta frequency during higher cognitive control loads. Simulated models show the slower theta frequency increases the probability of success in more difficult scenarios [40], better adapting to the task demands. Given the diverse roles of theta band activity and phase synchrony in executive tasks, it is not surprising that it is highly predictive of CEN activity across numerous executive tasks, as seen in Figure 4.

Theta band activity also plays a major role in the retrieval of autobiographical memories, increasing in amplitude during autobiographical memory recall [18]. It also provides a mechanism for a vmPFC-linked schema instantiation model, suggesting that the vmPFC modulates more posterior long-term memory representations [15]. Additionally, cross-frequency theta-gamma coupling between medial temporal lobe (MTL) structures and cortical regions is critical for the recall of autobiographical memories, and is disrupted in individuals with severely deficient autobiographical memory (SDAM) [13]. Our findings align broadly with this large body of literature, with theta-gamma cross-frequency coupling predicting DMN activity (see Figure 4), further asserting that the trained classifiers identified critical EEG features that represent processes inherent to each ICN.

Interestingly, cross-frequency coupling between delta, alpha and gamma bands was found to be predictive of SN activity, with the gamma and delta bands being concentrated at the right frontal electrodes (see Figure 4i). Combined with the predominantly right lateralization of theta-gamma cross-frequency coupling indicative of DMN activity (see Figure 4h), our results point towards a broadband salience hub anchored close to the right frontal region, which is also active during DMN-linked tasks such as autobiographical memory recall and spatial navigation. This pattern of activity is consistent with activation of the right anterior insula (rAI), a major hub of the salience network [25]. In the context of the tri-network model, the SN is thought to control the switch between the CEN and DMN, and is found to co-activate with the task-appropriate network during the task used in Cohort A [44]. As such, across a wider range of tasks, this node might need to synchronize with numerous brain regions across a wide range of frequencies, as seen in the cross-frequency results of Figure 4i.

In sum, the data-driven analysis pipeline used in this study identified EEG features that captured the core oscillatory dynamics of critical CEN, DMN and SN functions. Furthermore, the classification results demonstrate the collective utility of these salient features in discriminating between CEN, DMN and SN activity.

While this study focused on the three major ICNs within the tri-network model, the discussed approach is equally applicable to identifying unique EEG signatures of other fMRI-derived ICNs, such as ventral and dorsal attention networks (VAN/DAN), somato-sensory networks and the motor network [52, 45]. This opens up the possibility of using EEG to detect the ICN dynamics of any fMRI-derived ICNs, greatly improving the accessibility of such measures. To this end, the code developed for this analysis pipeline, along with a pretrained generalized classifier, is available for download from github.com/saurabhshaw/EEGnet.

The approach used in this study estimated ICN features using connectivity measures of electrode-level activity, rather than the traditionally used, and much more computationally complex, source-localization techniques. This permitted the exploration of a massive (4M-dimensional) feature space to find optimal features predictive of ICN activity. Unfortunately, when working with features in electrode space (rather than source space), there is the problem of volume conduction, which distorts the neural signal measured at each electrode. To mitigate this, phase-based functional connectivity measures such as phase lag index (PLI) and directed phase lag index (dPLI) were included in the feature space, given their immunity to volume conduction-based distortions [19].

Furthermore, staying in electrode-space, rather than transforming the data to source space, also reduced the computational time required to extract relevant features and predict the predominantly active ICN of a new EEG window. Further algorithmic optimizations were made to parallelize the computation of these features, allowing the use of massively parallel super-computing clusters to accelerate these computations. Future work can use this ability to make rapid predictions of ICN activity to develop novel network-based neurofeedback interventions to directly target the network dysfunction observed in numerous psychopathologies.

In conclusion, this paper takes an important step towards enabling EEG-based investigation of ICN dynamics, greatly increasing the accessibility of such measures in scenarios where fMRI-based ICN assessment might not be practical or economically feasible. This is particularly relevant for studying and characterizing complex psychopathologies such as PTSD, with various disease subtypes that require drastically different treatment plans. For example, the DMN is erroneously recruited instead of the CEN during working memory in PTSD patients [8], and the pattern of resting state ICN activity can be predictive of the dissociative PTSD subtype [30]. Hence, identifying the pattern of disrupted ICN switching dynamics can greatly help in characterizing the patients’ psychopathology and inform their treatment plans. This study makes it clinically feasible to do so by successfully applying cutting-edge machine learning techniques, such as semi-supervised learning, to create an analysis pipeline capable of detecting ICN activation using EEG data alone with high accuracy.

## 4. Methods

In this study, we developed a purely EEG-based model capable of classifying and tracking ICNs, traditionally investigated using fMRI. To accomplish this, we used simultaneously acquired EEG-fMRI data to identify an optimal EEG-based feature set that could predict the fMRI-derived ICN activation. To maximize the ability of our model to predict ICN activity across a wide range of cognitive tasks, we collected simultaneous EEG-fMRI data from two cohorts of participants that engaged in two different task-switching paradigms, as shown in Figure 1A and B.

### 4.1. Cohort A Task

The first cohort consisted of 14 participants that dynamically switched between a 2-back working memory (WM) task and an autobiographical memory (ABM) retrieval task, designed to activate the CEN, DMN respectively, and at task-switching points, to activate the SN.

Prior to scanning, each participant recorded up to 10 positive or neutral autobiographical memories in vivid detail, as well as descriptive words corresponding to each memory, that would serve as retrieval cues during the ABM task. Following this, they completed a 1 hour 20 min long memory assessment in the MRI scanner, comprised of randomly ordered 30-second blocks of either cued autobiographical memory retrieval or a 2-back working memory task. Each ABM block included 10 cues pertaining to one of the previously recorded memories, while each WM block included a sequence of 10 to-be-remembered, commonly used English words, each shown for 2 seconds. The participants were instructed to recall the cued memory shown during the ABM blocks, and to remember the word shown two words ago during WM blocks. To assess whether the participant was staying on task, at random points within each block, they were asked to perform a 4-alternative forced choice (4-AFC) trial. During ABM blocks, the 4-AFC trial required the participant to select, from four displayed words, the one representing the memory they were currently recalling from a selection of four words. During WM blocks, the 4-AFC trial required the participant to choose, from four displayed words, the one they saw two words ago. Each pending task switch was cued with 2-second trial showing either Word Memory or Autobiographical Memory, representing an upcoming WM block or ABM block respectively. Each run of sixteen blocks was followed by a 60 second rest period. Participants completed as many 16-block runs as possible within 80 minutes, up to a maximum of 4 runs (64 blocks - 32 ABM, 32 WM). The two distinct block types (ABM and WM) were predicted to activate the DMN and CEN respectively, while the cue to a pending task switch between blocks was expected to activate the SN. Shaw *et al*. [44] reports the activation dynamics of the DMN, CEN and SN observed during this study, using fMRI data alone.

### 4.2. Cohort B Task

The second cohort consisted of 12 participants that dynamically switched between 7 different tasks. Each participant performed a total of 18 trials of each of the following 7 tasks. This was split into three runs of 6 trials of each of the 7 different task types (42 trials/run), ensuring that trials of the same task type were not repeated more than twice in a row. Each trial began with a pre-recorded, single-word auditory cue indicating the type of the upcoming trial to the participant, followed by a 16-second imagery period. The participant was shown a fixation cross at the center of the screen during this imagery period to avoid any eye movements.

1. **Sport-related motor imagery** Participants were asked to imagine intensely performing a sport or full-body activity (*e.g*., dancing, jumping jacks) of their choice. Additionally, they were instructed to focus on the kinesthetic and somatosensory aspects of that activity rather than on visual aspects.
2. **Navigation imagery** Participants were asked to imagine navigating around their home from room to room, paying attention to all aspects of the room (*e.g*., placement of furniture, decor, objects in room).
3. **Music imagery** Participants were asked to imagine listening to a familiar song of their choice through headphones, while concentrating on all aspects of the song (e.g. the melody, instrumentation).
4. **Mental arithmetic** Participants were asked to count backwards by threes from a random 3-digit number of their choice. They were instructed to choose a different 3-digit number for each trial.
5. **Finger tapping imagery** Participants were asked to imagine pushing a button with each of the fingers of the right hand in succession, repeatedly, focusing on the somatosensory and kinesthetic rather than visual aspects of the imagery.
6. **Running imagery** Similar to the sport imagery condition, the participants were instructed to imagine only running, while attending to the kinesthetic and somatosensory aspects of the imagery.
7. **Rest** Participants were asked to clear their mind and think of nothing in particular.

Of these seven tasks, only two tasks were expected to maximally activate the DMN, *i.e*. navigation imagery and rest. The rest of the tasks were expected to activate a combination of attention and somatosensory ICNs that would work in tandem to successfully perform the relevant imagery. Other results exploring the strength of mental imagery in this dataset have been previously reported in Harrison *et al*. [16].

### 4.3. Data acquisition

Data acquisition for both cohorts was performed at the same site, using the same EEG system and MRI machine.

All EEG data were acquired using a BrainProducts (Brain Products GmbH, Gilching, Germany) 64 channel MR compatible EEG cap, at a sampling rate of 5000Hz. The electrode locations followed the extended international 10-20 system of electrode placement, with the reference at FCz and the ground at AFz. The impedance of each electrode was kept below 10kΩ. The physical setup used for this acquisition is further described in Shaw [42].

All MRI data were acquired using a GE Discovery MR750 3T MRI scanner and an 8 channel RF coil (General Electric Healthcare, Milwaukee, WI). A high resolution anatomical scan was acquired for each participant using an IR prepped axial 3D FSPGR sequence (*Cohort A*: TI/TR/TE=450/7.7/2.2ms, 12° flip angle, 240mm FOV, 2mm-thick slices of size 320 × 192, interpolated to 512 × 512; *Cohort B*: TI/TR/TE=900/10.312/3.22ms, 9° flip angle, 240 × 180mm FOV, 1mm-thick slices of size 512 × 248, interpolated to 512 × 512). These individualized anatomical scans were used to prescribe the fMRI scans, acquired using a 2D GRE EPI sequence (*Cohort A*: TR/TE=2000/35ms, 90° flip angle, 240mm FOV, 3.8mm-thick slices of size 64 × 64, 39 slices/volume interleaved, 300 volumes per functional run; *Cohort B*: TR/TE=3200/35ms, 90° flip angle, 240mm FOV, 4mm-thick slices of size 64 × 64, 40 slices/volume interleaved, 214 volumes per functional run).

### 4.4. Pre-processing

#### 4.4.1. fMRI pre-processing

All MRI pre-processing steps were performed using SPM12 and the CONN toolbox [58]. The fMRI scans were first realigned and unwarped, followed by motion correction, performed by adding the participant’s estimated motion (6 DOF) as a first-level covariate in a denoising general linear model (GLM). This was followed by frequency-domain phase shift slice timing correction (STC), and ART-based identification of outlier scans to be scrubbed. The functional scans were then normalized to the MNI152 atlas by aligning them to each participant’s MNI-aligned anatomical scan, followed by segmentation of the functional scans to remove skull, white matter and cerebral spinal fluid (CSF). Spatial smoothing was applied by convolving the BOLD signal with a 2mm Gaussian kernel. Finally, the BOLD data was band-pass filtered between 0.008Hz–0.09Hz.

#### 4.4.2. EEG pre-processing

The gradient artefact (GA) in the EEG data, collected concurrently with the fMRI data, was filtered using a parallel optimized version of the FASTR gradient artefact removal toolbox Shaw [42], which relies on the GA template subtraction algorithm [1]. This was followed by the detection of the QRS complex using data from the ECG lead, which was then used to filter the ballistocardiogram (BCG) using optimal basis set filtering (OBS). The artefact-free EEG data were then downsampled from 5000Hz to 500Hz, followed by temporal band-pass filtering into six different frequency bands - full (1-50Hz), delta (1-4Hz), theta (4-8Hz), alpha (8-13Hz), beta (13-30Hz), and gamma (30-50Hz). All temporal filtering was performed twice, once in the forward, and once in the reverse direction for zero phase lag using a 12-order Butterworth IIR filter. Bad channels were then detected and removed based on their spectral characteristics, followed by an ICA decomposition using the FAST-ICA algorithm implemented within EEGLAB [9]. The EEG components representing ocular artefacts, eye blinks and muscle artefacts were detected and removed using the automated ICLabel tool. Finally, the EEG data were referenced to average EEG channel data, after which the previously removed bad channels were interpolated using spherical interpolation. The bad channels were only interpolated if the number of bad channels was less than 5% of the total number of channels, while also ensuring no two bad channels were neighbours.

### 4.5. Feature Extraction

#### 4.5.1. FMRI Independent Component Analysis (ICA)

A group independent component analysis (group-ICA) [3] was performed on the denoised fMRI voxel-level data using the iterative FastICA algorithm. This analysis identified 20 mutually independent spatiotemporal patterns of activity, known to represent ICNs. The spatial overlaps (Dice coefficients) of the group-ICA components with known ICNs[45] were used to label the ICA components, identifying the components that corresponded to CEN, DMN or SN sub-networks, as shown in Figure 1C.iii. These group-level ICA components were then back-projected to individual participants’ data using GICA back-projection [11], to obtain the activity timeseries of each ICA component. The activity of the CEN, DMN and SN networks were identified by averaging the activity of their constituent subnetworks. Using a sliding-window method, the time courses were segmented into overlapping windows of length 10 seconds (5 TRs) for cohort A and 9.6 seconds (3 TRs) for cohort B. Using a time step of 1 fMRI time point, the time series were segmented into 1184 windows for each participant in cohort A and 636 windows for each participant in cohort B. Each window was labeled with the most active ICN by first thresholding the activity at 70%, followed by comparing the activity of the CEN, DMN and SN during that time period to pick the ICN with the highest activity (shown in Figure 1C.v). These labels were used as the “ground-truth” labels of the ICN activity for all subsequent analyses.

#### 4.5.2. EEG Feature Extraction

A large battery of functional connectivity features (>40 million) were extracted from the EEG data corresponding to each fMRI window. These included single-frequency features representing connectivity between two channels within the same frequency band: coherence (COH), phase lag index (PLI) and directed phase lag index (dPLI); and cross-channel cross-frequency coupling features representing connectivity between two channels, across two different frequency bands: the synchronization index (SI) and phase-amplitude coupling (PAC). Each single-frequency feature was computed for each of the 6 frequency bands (full band, delta, theta, alpha, beta, gamma) described above, and each cross-frequency feature was computed for all possible pairs of the 6 frequency bands. Each fMRI window was further divided into 99 windows of width 200ms, which were used to estimate the features, described in more detail below.

***Coherence (COH)***. represents the synchrony between two channels by comparing their power spectral densities [59], and is computed as follows

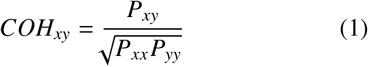

where *P_xx_* and *P_yy_* are the power spectral densities (PSD) of two channels *x* and *y*, and *P_xy_* is the cross-spectral density (Fourier Transform of crosscorrelation) of the signals *x* and *y*. *COH_xy_* values range between 0 and 1, with 0 representing no coherence between the two signals and 1 representing perfect coherence. Since the power spectral densities are heavily dependent on the signals’ amplitudes, COH is sensitive to volume conduction effects.

***Phase Lag Index (PLI)***. is another measure of functional connectivity that addresses the susceptibility of COH-based measures to volume conduction effects. PLI estimates the functional connectivity between two channels (*x* and *y*), by estimating the phase-synchronization between them. This relies on the assumption that two channels are functionally connected if there is a consistent phase delay in the signals coming from the two channels. This is defined in terms of their cross-spectrum, given by *C_xy_* = *H*(*x*) · *H*(*y*)*, where *H*(·) represents the Hilbert transform and * represents the complex conjugate. The PLI is then defined as *PLI_xy_* = |〈*sgn*(*Im*(*C_xy_*))〉|, where *Im*(*C_xy_*) is the imaginary part of the cross-spectrum (*C_xy_*), *sgn*(·) is the “sign” operator, and 〈·〉 is the expected value operator. It is important to note that *Im*(*C_xy_*) is equivalent to Δ*ϕ_xy_*-based definition in Stam *et al*. [50]. To minimize spurious noise, we used the weighted PLI variant in this study [54], where the *sgn*(*Im*(*C_xy_*)) is weighted by the imaginary component of the cross-spectrum (|*Im*(*C_xy_*)|), as given in Equation 2.

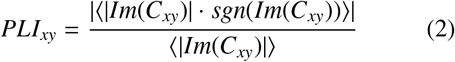

PLI values lie between 0 and 1, with 0 representing no consistent phase synchrony between channels *x* and *y*, and 1 representing perfect phase-locking. To minimize spurious PLI values, every PLI estimate was compared against its surrogate, estimated by the PLI of the signals spliced at random time points. Only those PLI values that were significantly different from their surrogates were accepted.

***Directed Phase Lag Index (dPLI)***. is a variant of PLI that retains information on phase directionality [49]. This relies on the assumption that lagging signals occur downstream from leading signals, establishing a directional link going from the leading channel to the lagging channel. Equation 3 was used to compute the dPLI for signals from two channels, *x* and *y*, of length *N*.

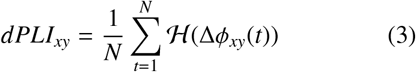

where Δ*ϕ_xy_*(*t*) = *ϕ_x_*(*t*) – *ϕ_y_*(*t*) is the difference in phases of the two signals, and 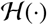 is the Heaviside step function. The value of *dPLI_xy_* represents the direction of functional connectivity, with 0.5 < *dPLI_xy_* ≤ 1 representing *x* leading *y* (*x* → *y*), and 0 ≤ *dPLI_xy_* < 0.5 representing *x* lagging *y* (*y* → *x*). Given its similarity, it is not suprising that PLI and dPLI are related: *PLI_xy_* = 2|0.5 – *dPLI_xy_*|. Similar to PLI estimates, only the dPLI estimates significantly different from its surrogates were retained.

***Phase Amplitude Coupling (PAC)***. is a measure of phase-amplitude cross-frequency coupling between two channels, specifically probing the impact of the phase of the signal in the lower frequency band on the amplitude of the signal in the higher frequency band. Among the several methods of estimating PAC, the modulation index (MI) approach was used due to its superior performance [53]. This approach estimates the PAC between two channels *x* and *y*, between two frequency bands *f_A_* and *f_B_* (*f_A_* < *f_B_*) using the following steps:

1. Filter the *x* and *y* channel data into the respective frequency bands, creating *x_f_A__* and *y_f_B__*.
2. Extract the phase time series of the lower frequency signal using: 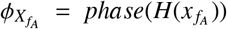, and the amplitude time series of the higher frequency signal using: 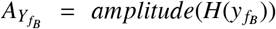, where *H*(·) is the Hilbert transform.
3. Bin the lower frequency phases in 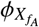 into *N* bins: 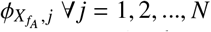, and find the mean higher frequency amplitudes for each phase bin *j*, denoted by 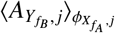. This amplitude distribution is normalized by its sum over all bins, as follows

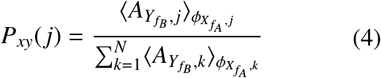 In the case of no PAC, the amplitude distribution *P_xy_*(*j*) is expected to be flat (uniform distribution) since the higher frequency amplitudes will not vary with the lower frequency phase. *N* = 20 was used for estimating the amplitude distribution.
4. The phase amplitude coupling is then estimated as the Kullback-Liebler (KL) divergence (*D_KL_*(·)) between the observed amplitude distribution *P_xy_*(*j*) and the uniform distribution *U*, normalized by the maximal possible entropy value (occurs for the uniform distribution).

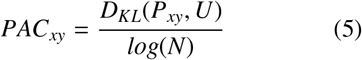

This procedure was repeated for each channel-frequency band pair. Each PAC estimate was retained only if it was significantly different from its surrogate, estimated in a manner similar to the PLI and dPLI measures.

***Synchronization Index (SI)***. is another measure of cross-frequency coupling between two channels, probing the impact of the phase of the signal in the lower frequency band on the phase of the power time series of the signal in the higher frequency band [6]. The following steps were followed to estimate the SI between two channels *x* and *y*, between two frequency bands *f_A_* and *f_B_*(*f_A_* < *f_B_*).

1. Filter the *x* and *y* channel data into the higher frequency band, creating *y_f_B__*.
2. Extract the power time series of the signal in the higher frequency band using: 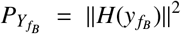, where *H*(·) is the Hilbert transform. To identify rhythmic fluctuations in this power time series at the lower frequency, compute the FFT of 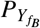 within the lower frequency band (*f_A_*). The peak in this FFT 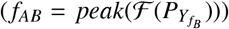 is the synchronization frequency within *f_A_*, which is then used to fine tune the bounds of *f_A_*. The revised lower frequency band 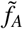 is defined by picking a window around the empirically identified synchronization frequency (*f_AB_*), using 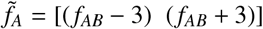.
3. Extract the phase time series of the signal within this revised lower frequency band 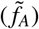 using: 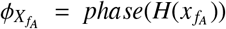, and the phase time series of the higher frequency band power time series using: 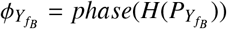.
4. The synchronization index is then calculated using Equation 6

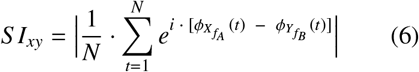

This procedure was repeated for each channel-frequency band pair. Each SI estimate was retained only if it was significantly different from its surrogate, estimated in a manner similar to the PAC, PLI and dPLI measures.

### 4.6. Feature selection

The feature set was reduced from > 40million features to 5000 features by using a hierarchical version of the popular minimum redundancy maximum relevance (mRMR) feature selection algorithm. mRMR minimizes the mutual information between features while maximizing the mutual information between the features and the class labels [33]. The hierarchical mRMR consisted of repeatedly applying mRMR to identify the top 1000 features at the channel level, followed by selection of the top 1000 features at the frequency band level, followed by selection of the top 1000 features at the window level. This resulted in 1000 top features for each of the five feature types described above, which were then concatenated to create the final 5000-dimensional feature set used for classification. This process is illustrated in Figure 1C.vi.

### 4.7. Classification

A multi-class support vector machine was trained on the three-way classification task of predicting the most active ICN during each fMRI window, using only the 5000-dimensional EEG feature set identified in section 4.6. Three classifiers were trained for each participant using the identified feature set: a generalized classifier, a semi-supervised individualized classifier and a fully-supervised individualized classifier. The generalized classifier was trained using data from all participants, except the participant being tested (leave-one-out cross validation), followed by prediction of the class labels of the test-participant. The semi-supervised individualized classifier used the highly confident label predictions as “expert” labels, while leaving the rest of the time windows unlabelled to select a participant-specific feature set, and use it to classify the unlabelled time points. Finally, the fully-supervised individualized classifier was trained on a subset of the data from the same participant being tested, ensuring that the training set and test set did not contain overlapping windows. A 75/25 train/test split was used with 20-fold cross validation for testing the performance of this classifier.

